# How low can you go: sex identification from low-quantity sequencing data despite lacking assembled sex chromosomes

**DOI:** 10.1101/2021.11.04.467120

**Authors:** Andrea A. Cabrera, Alba Rey-Iglesia, Marie Louis, Mikkel Skovrind, Michael V Westbury, Eline D Lorenzen

**Affiliations:** Globe Institute, University of Copenhagen, Øster Voldgade 5-7, 1350 Copenhagen K, Denmark; Greenland Institute of Natural Resources, Kivioq 2, Nuuk 3900, Greenland

**Keywords:** Low-coverage, sex assessment, molecular sexing, sex chromosome, bioinformatics

## Abstract

Accurate sex identification is crucial for elucidating the biology of a species. In the absence of directly observable sexual characteristics, sex identification of wild fauna can be challenging, if not impossible. Molecular sexing offers a powerful alternative to morphological sexing approaches. Here, we present SeXY, a novel sex-identification pipeline, for very low-coverage shotgun sequencing data from a single individual. SeXY was designed to utilise low-effort screening data for sex identification and does not require a conspecific sex-chromosome assembly as reference. We assess the accuracy of our pipeline to data quantity by downsampling sequencing data from 100,000 to 1,000 mapped reads, and to reference genome selection by mapping to a variety of reference genomes of various qualities and phylogenetic distance. We show that our method is 100% accurate when mapping to a high-quality (highly contiguous N50 > 30 Mb) conspecific genome, even down to 1,000 mapped reads. For lower-quality reference assemblies (N50 < 30 Mb), our method is 100% accurate with 50,000 mapped reads, regardless of reference assembly quality or phylogenetic distance. The SeXY pipeline provides several advantages over previously implemented methods; SeXY (i) requires sequencing data from only a single individual, (ii) does not require assembled conspecific sex-chromosomes, or even a conspecific reference assembly, (iii) takes into account variation in coverage across the genome, and (iv) is accurate with only 1,000 mapped reads in many cases.

## Introduction

Accurate sex identification is critical for elucidating the life history, behaviour, social structure, and demography of a species. It is particularly important for taxa where females and males differ in prey preference (e.g., Louis et al., 2021), social interactions and mating behaviour (e.g., Amos et al., 1993; Pečnerová et al., 2017), and seasonal movements and dispersal (e.g., Dobson & Stephen Dobson, 1982; Gower et al., 2019; Greenwood, 1980). Reliable sex identification may also help to elucidate the impacts of past and present anthropogenic activities on wildlife, including prehistoric hunting or domestication practices (e.g., Nistelberger et al., 2019), and the identification of the sex of and sex biases in ongoing wildlife poaching (e.g., Malisa et al., 2005).

In the absence of directly observable sexual characteristics, such as morphology or behaviour (Fairbairn et al., 2008), sex identification of wild fauna remains challenging, if not impossible. An additional challenge for research based on museum or palaeontological specimens is the sex identification of skeletal remains. In most cases, such as in the (sub-)fossil record, only small skeletal fragments are available. Osteological sex determination may also be limited by the degree of preservation, the age of the individual, or access to appropriate reference material with which to compare (Buonasera et al., 2020).

Molecular sexing can be used as an alternative to morphological sexing; it only requires a small tissue sample (Hrovatin & Kunej, 2018), and may even be applied to environmental samples (e.g., Durnin et al., 2007). Many molecular sexing techniques utilise information regarding the homogametic and heterogametic sexes. In mammals, and in many fishes, females are homogametic and males are heterogametic with XX and XY chromosomes, respectively (Ellegren, 2000; Í Kongsstovu et al., 2020; Moore, 1925). In birds and certain reptiles, the pattern is reversed, with females having ZW and males having ZZ chromosomes.

For tissue samples with high-quality DNA, molecular sex identification is relatively fast, inexpensive, and straightforward. Methods for mammals include PCR-based techniques that (i) amplify the SRY gene of the Y chromosome (Bryja & Konečný, 2003; Pomp et al., 1995), or (ii) target specific regions of the ZFX and ZFY genes found on the X and Y chromosomes, respectively (Aasen & Medrano, 1990; e.g., Bérubé & Palsbøll, 1996; Curtis et al., 2007). However, these approaches require specific laboratory work targeting loci in sex chromosomes (e.g., Ahlering et al., 2011), and are not suitable for samples with highly fragmented and/or degraded DNA, such as material not specifically sampled and preserved for DNA analysis (including skeletal remains, wildlife products, and museum specimens). PCR failure in method (i) and a biased amplification of the ZFX over the ZFY region (Sinding et al., 2016) in method (ii) may cause males to be misidentified as females.

The analysis of shotgun sequencing data offers a more robust approach to identify the sex of an individual; endogenous shotgun data can be retrieved from samples with low-quality DNA, with no additional laboratory procedures required to specifically target loci on sex chromosomes. Sex-identification pipelines for DNA data with a low number of target reads were originally developed for human ancient DNA data, and were based on either the ratio of number of reads aligning to the X and Y chromosomes (Skoglund et al., 2013), or on the ratio of number of reads aligning to the X chromosome *versus* the autosomes (Mittnik et al., 2016). This last method has recently been utilised on elephants and other mammalian taxa for which the X chromosome of either a conspecific or a related reference genome is available (Bro-Jørgensen et al., 2021; de Flamingh et al., 2020). Although this approach has been shown to be efficient down to ∼10,000 mapped sequencing reads, it requires either a conspecific chromosome-level assembly with known sex chromosomes, or mapping to a more distantly related chromosome-level assembly, with decreased mapping efficiency as a result.

Reference genome assemblies from non-model vertebrate species with assembled sex chromosomes are relatively scarce. Available mammalian genome assemblies with at least one sex chromosome (most commonly the X chromosome) include humans, several domesticates such as cat (*Felis catus*), cow (*Bos taurus*), dog (*Canis familiaris*), horse (*Equus caballus*), sheep (*Ovis aries*), and wild species such as blue whale (*Balaenoptera musculus*), bottlenose dolphin (*Tursiops truncatus*), greater horseshoe bat (*Rhinolophus ferrumequinum*), gorilla (*Gorilla gorilla*), meerkat (*Suricata suricatta*), orangutan (*Pongo pygmaeus*), and vaquita (*Phocoena sinus*) (Cabrera et al., 2021; de Flamingh et al., 2020). In the absence of a conspecific chromosome-level assembly, alternative approaches can be used to identify scaffolds originating from sex chromosomes. Approaches include synteny-based, whole-genome alignments (e.g., Grabherr et al., 2010), and the estimation of relative coverage of each scaffold using data from known females and males of the target species (reviewed in Palmer et al., 2019). Sex identification using synteny or coverage approaches has been applied in some studies using ancient (e.g., Kirch et al., 2021) or degraded DNA (e.g., Skovrind et al., 2019). However, the pipelines have been developed for specific species and datasets, and an assessment of the minimum level of required sequencing data and of the impact of reference genome assembly choice is lacking.

Methods exist that circumvent the need to *a priori* identify sex-linked scaffolds. For example, a recent fast and automated method “Sex Assignment Through Coverage” uses principal component analysis to identify sex-related scaffolds, and the sex of an individual (Nursyifa et al., 2021). This approach holds promise for studies that include a relatively large number of samples, as the method requires a set of both male and female samples. However, these sample requirements may not always be met.

Here, we present a sex-identification method (SeXY) for taxa lacking a conspecific chromosome-level assembly. The method can be applied to shotgun sequencing data from mammals, and potentially to any species with a heterogametic sex (e.g, birds, and some reptiles, fish, and insects) in which the target and reference species share the same sex-determination system (i.e., same sex chromosomes, same sex determining locus, same sex determining gene). We use a synteny-based approach to identify putative X-linked scaffolds in the reference assembly, and determine sex using the expectation that males (in mammals) have half the amount of X-chromosome genetic material compared to females. We assessed the robustness of this method using raw shotgun sequencing data from two target marine mammal species: beluga whale (*Delphinapterus leucas*) and polar bear (*Ursus maritimus*). The read data were subsampled and mapped to reference assemblies of various quality and phylogenetic distance. We show our approach to be highly accurate (i) with as few as 1,000 mapped reads when mapping to a high-quality (chromosome level) reference genome assembly, or as few as 50,000 mapped reads when mapping to a lower-quality reference genome assembly (N50 < 30 million base pairs (Mb)); (ii) also when using a phylogenetically distant reference genome assembly; and (iii) without known sex chromosomes.

## Materials and Methods

The SeXY method requires (i) raw shotgun sequencing reads of a target individual;(ii) an assembled genome from either a conspecific or related species (RefGEN) with the same sex determination system; and (iii) assembled X and Y chromosomes (RefX and RefY, respectively), which can be either from the same or another species than the RefGEN.

We assessed the applicability of SeXY using data from two target species: beluga and polar bear. We also assessed the impact of reference assembly using four RefGEN of varying quality and phylogenetic distance to each target species, and two reference sex chromosome assemblies (each comprising RefX and RefY) from species of varying phylogenetic distance. To ascertain the applicability of our method to specimens with low DNA yield, we additionally tested the impact of the number of mapped reads on the sex determination using various downsamplings ranging from 100,000 to 1,000 mapped reads.

### 1. Target species data and reference assemblies

We used publicly available Illumina shotgun sequencing reads from ten beluga and ten polar bear individuals (Supplementary Table S1). Each species dataset comprised five females and five males. As we were interested in results produced with <=100,000 mapped reads only, all read files were randomly downsampled to one million reads using the sample option in seqtk v1.3 (https://github.com/lh3/seqtk), to reduce computational time during the mapping step.

To evaluate the impact of reference genome assembly, we used four reference assemblies (RefGEN) for each target species (beluga, polar bear): two conspecific RefGEN of differing assembly quality, and two RefGEN from more divergent species (Figure 2; Supplementary Table 2). To reduce computational time and memory usage, all scaffolds < 10 kilobase (kb) were removed from the RefGEN files and excluded from downstream analyses using reformat.sh from the BBmap toolsuite (Bushnell, 2014).

For beluga, we included two beluga reference assemblies: one of lower quality (non-chromosome-level) (Beluga v1, N50 161 kb (Jones et al., 2017)) and one highly contiguous (non-chromosome-level) (Beluga v3, N50 31 Mb (Dudchenko et al., 2017, 2018)). We also included a relatively low quality killer whale (*Orcinus orca*) assembly (Orca, N50 13 Mb (Foote et al., 2015)) and a chromosome-level cow assembly (Cow, N50 103 Mb (Zimin et al., 2009)). Assuming a divergence time between the beluga and killer whale of ∼19 million years ago (Ma) (McGowen et al., 2020) and a yearly mutation rate for beluga of 5.16 × 10^−10^ (Westbury et al., 2019), the divergence between the beluga and killer whale genomes is estimated at ∼2%. The divergence between the beluga and cow genomes is estimated at ∼6.8% assuming a divergence time of ∼66 Ma (McGowen et al., 2020), and above-mentioned beluga mutation rate.

For polar bear, we included two polar bear reference assemblies: the lower quality Polar bear v1, N50 16 Mb (Liu et al., 2014) and the chromosome-level Polar bear v1 HiC, N50 71 Mb (Dudchenko et al., 2017, 2018). We also included a chromosome-level panda (*Ailuropoda melanoleuc*a) assembly (Panda, N50 129 Mb (Fan et al., 2019)), and a chromosome-level dog assembly (Dog, N50 64 Mb (Lindblad-Toh et al., 2005)). The estimated divergence between the polar bear and panda genomes is ∼6.4%, assuming a divergence time of ∼19.5 Ma (Hu et al., 2017) and a mutation rate for polar bear of 1.6 × 10^−9^ (Liu et al., 2014). The divergence between the polar bear and dog genomes is estimated at ∼17%, assuming a divergence time of ∼52 Ma (Hu et al., 2017) and above-mentioned polar bear mutation rate.

### 2. Identification of putative sex-linked and autosomal scaffolds

We identified scaffolds putatively originating from sex chromosomes (both X and Y) from all RefGEN lacking assembled sex chromosomes as well as from Cow and Dog, which include assembled sex chromosomes. We did this by aligning each RefGEN with a designated pair of RefX and RefY assemblies, using satsuma synteny v2.1 (Grabherr et al., 2010) with default parameters (Figure 1). To increase efficiency and only run the synteny analysis once, we concatenated the RefX and RefY assemblies in one file.

**Figure 1.**
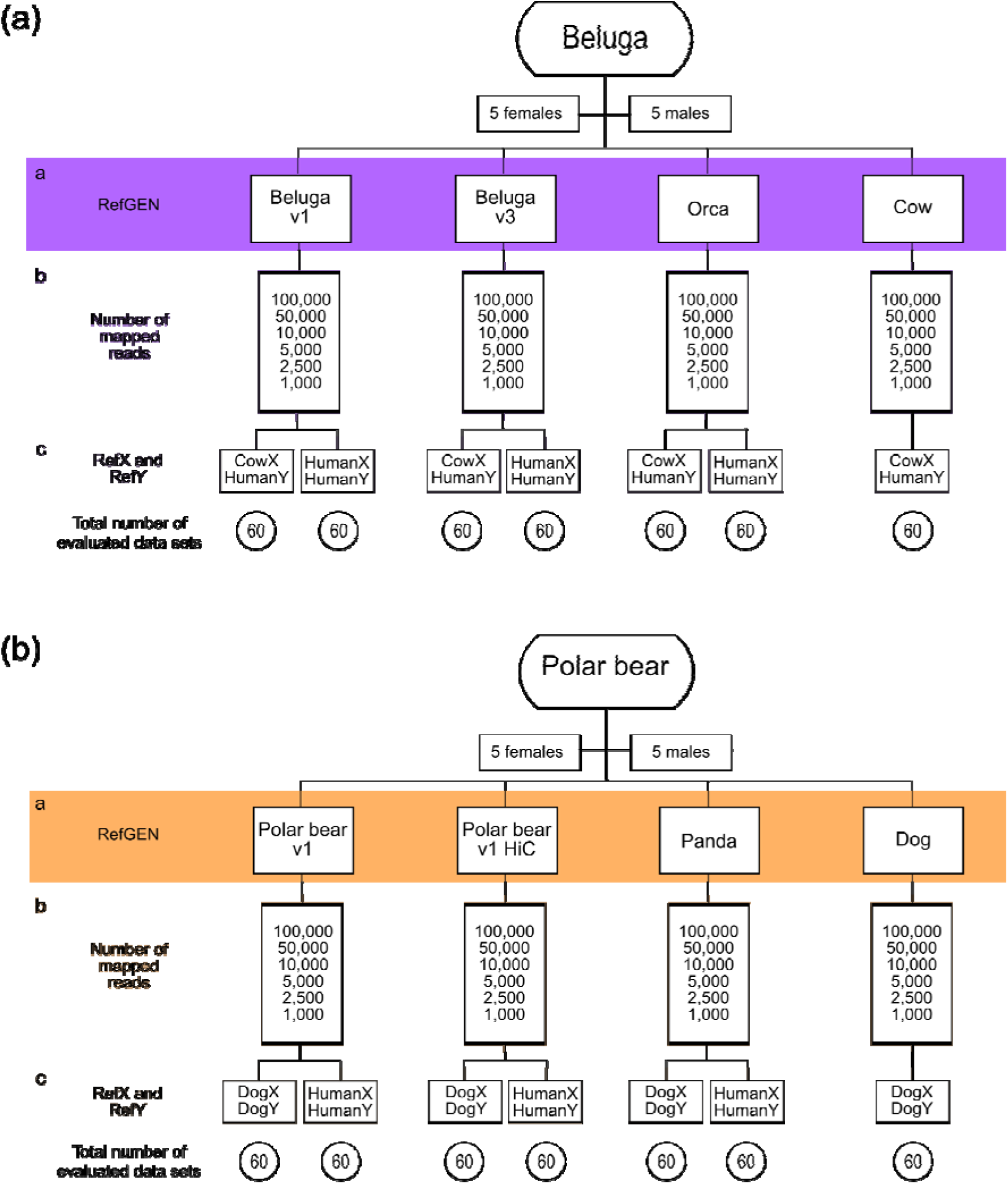
Schematic representation of the data sets and reference assemblies (RefGEN, RefX, RefY) analyzed for the two target species: beluga and polar bear. Each branch of the flowchart shows the evaluated combination of (a) reference genome assembly (RefGEN) used as mapping reference for the raw reads of each target species, (b) number of mapped reads of the target species (representing six independent data sets), and (c) reference sex-chromosome assembly (RefX and RefY) used to localize the sex-linked scaffolds (synteny). Total number of evaluated data sets per branch of the flow chart is shown at the bottom of the figure.

Although our method relies on comparing X chromosome and autosomal coverage (which we term X:A ratio), we included the Y chromosome to remove possible biases due to pseudoautosomal regions (homologous regions between the X and Y chromosomes) (Helena Mangs and Morris 2007). To reduce this bias, we removed any overlapping coordinates between the X-and Y-linked scaffold bed files using bedtools v.2.29.0 intersect (Quinlan & Hall, 2010). We identified putative autosomal scaffolds by removing the previously identified putative sex-linked scaffolds from each RefGEN.

We selected three RefX and RefY combinations: (i) HumanX and HumanY, (ii) CowX and HumanY, and (iii) DogX and DogY (Supplementary Table S3). The human sex chromosome assemblies were selected as they are the most well-assembled mammalian sex chromosomes available. We selected the cow and dog sex-chromosome assemblies, as they each represent the highest-quality, chromosome-level assemblies with defined sex chromosomes within the same phylogenetic order as each of our target species: beluga (Artiodactyla) and polar bear (Carnivora). For the cow, we used HumanY as there was no cow Y-chromosome available. We used the three RefX and RefY combinations to assess the influence of phylogenetic distance to the target species on downstream sex determination. For the cetacean/cow RefGEN dataset used for beluga, combinations (i) and (ii) were used (Figure 1a). For the bear/dog RefGEN dataset used for polar bear, combinations (i) and (iii) were used (Figure 1b) (Supplementary Table 3). For the Cow and Dog RefGENs only one combination of RefX and RefY was tested (CowX and HumanY for the former, and DogX and DogY for the latter). The estimated divergence between the beluga and human genomes is ∼9.9%, assuming a divergence time of ∼96 Ma (Kumar et al., 2017) and above-mentioned mutation rate for beluga. The divergence between the polar bear and human genomes is estimated at ∼31.4%, assuming a divergence time of ∼96 Ma (Kumar et al., 2017) and above-mentioned polar bear mutation rate.

### 3. Mapping and downsampling of mapped reads

Processing and mapping of raw beluga and polar bear sequencing reads to each designated RefGEN (Figure 1A) was performed using the Paleomix pipeline v.1.3.2 (Schubert et al., 2014). Adapter sequences were trimmed from the raw reads with AdapterRemoval v.2.3.1 (Schubert et al., 2014, 2016) using default settings and a minimum read length of 30 bp. Trimmed reads were mapped with BWA-MEM v.0.7.17 (Li, 2013) to each RefGEN. Mapped reads with mapping quality < 30 were removed using SAMtools v1.9 (Li et al., 2009). Duplicates were removed using Picard MarkDuplicates (http://broadinstitute.github.io/picard). The RefGENs used for mapping include both the autosome- and sex-chromosome scaffolds and should not include the mitochondrial genome. In our case, only in the low quality assembly Beluga v1 was the information regarding the mitochondrial genome not specified. In case the information is not specified, or the mitochondrial genome is included in the RefGEN, it is possible to first map the reads to a mitochondrial genome and exclude those mapped reads.

To evaluate the impact of number of mapped reads on genetic sex determination, we randomly downsampled the bam files to 100,000; 50,000; 10,000; 5,000; 2,500 and 1,000 mapped reads (Figure 2) using BBMap (Bushnell, 2014). We evaluated the differences in the mapping efficiency to each RefGEN, measured as the number of raw reads required to obtain a specific number of mapped reads (Figure 2, Supplementary Figure S1, and Supplementary Table S4).

**Figure 2.**
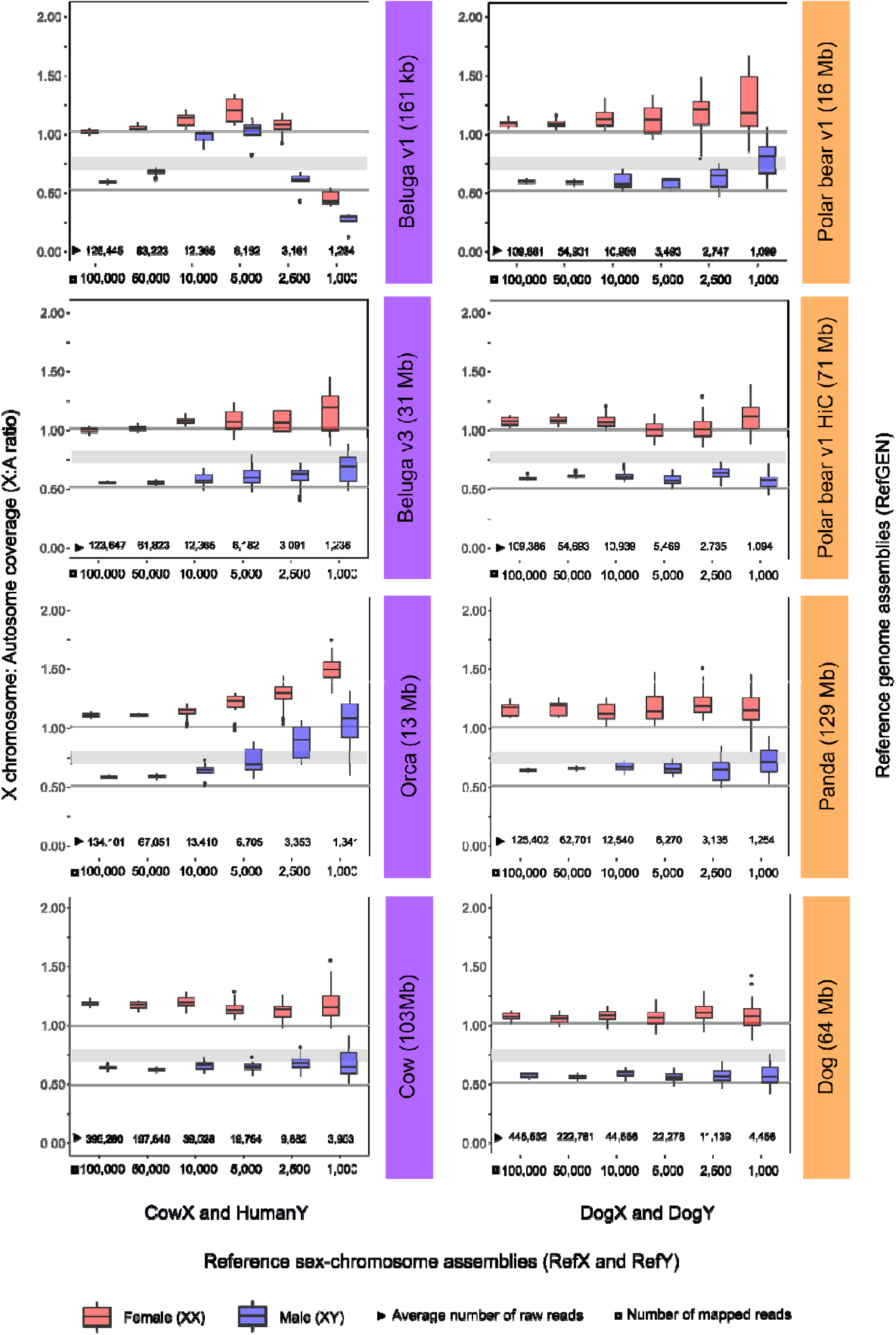
Sex determination of beluga and polar bear individuals using four reference genome assemblies (RefGEN), one combination of reference sex-chromosome assembly (RefX and RefY) for each target species, and various numbers of mapped reads. The ten beluga and ten polar bear individuals tested both comprised five females (red) and five males (blue). X axis shows number of mapped reads (square) and average number of raw reads necessary to obtain the required number of mapped reads (triangle). Y axis shows comparison of X chromosome and autosome coverage (X:A ratio) for each combination of RefGEN, RefX and RefY (CowX and HumanY, DogX and DogY), and number of mapped reads. Individuals were determined as females if their X:A ratio was >=0.8, and as males if their X:A ratio was <=0.7. Grey shaded horizontal bars indicate an X:A ratio of 0.7-0.8, which we interpreted as undetermined sex.

### 4. Sex determination

The sex of each individual was estimated based on the X chromosome:autosome coverage ratio (X:A ratio). We calculated the read depth of all sites from the X-linked scaffolds and from the autosomal scaffolds using SAMtools depth v.1.9 (Li et al., 2009), specifying minimum base and mapping qualities of 25. To take into account variation across genomic regions, we randomly selected 10 million sites from both X-linked and autosomal scaffolds independently, calculated the average coverage for those sites, and calculated the X:A ratio from the average coverages. This step was repeated ten times (Supplementary Table S5). As female mammals have two copies of the X chromosome, and males carry only one copy, we expected X:A ratios of ∼1 and ∼0.5 for females and males, respectively. We determined a female as correctly identified if the mean X:A ratio of the ten replicates was >=0.8 and a male if the mean X:A ratio of the ten replicates was <= 0.7. We considered a X:A ratio of 0.7 - 0.8 as ‘undetermined’ sex as used in previous studies (de Flamingh et al., 2020; Mittnik et al., 2016)

When interpreting the accuracy of the method, we considered (i) correctly determined sex; (ii) ‘undetermined’ sex, (iii) incorrectly determined sex (Supplementary Table S6). We did this to indicate whether accuracy below 100% was due to individuals with undetermined sex (with a X:A ratio of 0.7-0.8), or due to with incorrectly determined sex, as the latter is more detrimental to biological inference than simply the inability to determine sex.

## Results

### 1. Mapping

In agreement with previous results (Prasad et al., 2021), we found a decline in mapping efficiency as phylogenetic distance to the RefGEN increased (Supplementary Table S4). For the beluga dataset, the average percentage of raw reads successfully mapping and passing filters were as follows: Beluga v1 - 81%, Beluga v3 - 82%, Orca - 75%, and Cow - 25%. For the polar bear dataset, the average percentage was: Polar bear v1 - 91%, Polar bear v1 HiC - 91%, Panda - 80%, and Dog - 24%.

### 2. Sex determination

We found the sexing approach implemented in SeXY provided 100% accuracy in sex determination across all combinations of reference genome assembly (RefGEN) and reference sex-chromosome assemby (RefX, RefY), when 100,000 and 50,000 mapped reads were available (Figure 2, Table 1, Supplementary Figure S1, Supplementary Table S6-S7). Moreover, 100% accuracy was observed for most trials involving lower numbers of mapped reads; 10,000 and 5,000. Clear exceptions could be seen when using Beluga v1 (N50 161 kb) and Orca (N50 13 Mb) as RefGEN in the beluga data set. Inaccuracies were especially prevalent when the low-quality Beluga v1 RefGEN (N50 161 kb) was used; we found a marked decline in accuracy when using ≤10,000 mapped reads, with sex determination accuracy in some cases equivalent to random chance (down to 50%) (Table 1).

**Table 1:** Summary table showing percentage of correct sex determination across tested combinations of reference genome assembly (RefGEN), reference sex-chromosome assembly (RefX and RefY), and number of mapped reads. Results are shown for the beluga data and the cetacean/cow RefGEN assemblies tested (left columns), and for the polar bear data and the bear/dog RefGEN assemblies tested (right columns). The value below each RefGEN indicates the assembly N50. For cells with two estimates, the left value indicates estimates including undetermined sex, and the right value indicates estimates excluding undetermined sex. Only one value is included if both estimates were the same. Percentages in each cell are based on ten sample individuals; five females and five males. Sex determination for each indvidual was calculated using the average value of ten replicates, assuming a threshold of <= 0.7: male; 0.7-0.8: undetermined sex; >= 0.8: female. Corresponding summary table for tests using HumanX and HumanY as RefX and RefY, respectively, is provided in Supplementary Table S7.

|  | Beluga |  |  |  | Polar bear |  |  |  |
| --- | --- | --- | --- | --- | --- | --- | --- | --- |
| Number of mapped reads | Beluga v1 | Beluga v3 | Orca | Cow | Polar bear v1 | Polar bear v1 HiC | Panda | Dog |
|  | 161 kb | 31 Mb | 13 Mb | 103 Mb | 16 Mb | 71 Mb | 129 Mb | 64 Mb |
|  | CowX and HumanY |  |  |  | DogX and DogY |  |  |  |
| 100,000 | 100 | 100 | 100 | 100 | 100 | 100 | 100 | 100 |
| 50,000 | 100 | 100 | 100 | 100 | 100 | 100 | 100 | 100 |
| 10,000 | 50 | 100 | 90/100 | 100 | 100 | 100 | 100 | 100 |
| 5,000 | 50/56 | 90/100 | 80/89 | 100 | 100 | 100 | 100 | 100 |
| 2,500 | 100 | 100 | 50/63 | 80/100 | 100 | 100 | 90/100 | 100 |
| 1,000 | 50 | 80/100 | 60 | 80/89 | 70 | 100 | 80/89 | 100 |

Taken together, our results showed scaffold contiguity of the RefGEN influences the accuracy of sex determination more than phylogenetic distance. Across all trials, we found the highest percentage of correctly identified sex was obtained with highly contiguous (Beluga v3) or chromosome-level (Polar bear v1 HiC, Panda, Dog) RefGEN, regardless of whether the RefGEN was from a conspecific or a more divergent species (Table 1, Figure 2).

For the beluga dataset and CowX and HumanY RefXY (table 1 and figure 1), we found 100% accuracy in sex determination down to 10,000 mapped reads when using the higher-quality Beluga v3 (N50 31 Mb) and Cow (N50 103 Mb) RefGENs (Table 1). When we decreased the number of mapped reads below 5,000, we obtained a 10%-20% decrease in accuracy, which resulted in some undetermined individuals. However, for the trials where we were able to determine sex, the sex was determined with 100% accuracy down to 1,000 and 2,500 mapped reads with Beluga v3 and Cow as RefGEN, respectively.

When analysing the polar bear dataset and DogX and DogY RefXY (table 1 and figure 1), we found 100% accuracy in sex determination down to 5,000 mapped reads for all RefGEN. Both polar bear RefGENs (Polar bear v1, Polar bear v1 HiC) produced similar sex determination accuracies (Table 1), with 100% accuracy down to 2,500 mapped reads.

However, when we decreased the number of mapped reads to 1,000, mapping to the less contiguous Polar bear v1 correctly determined the sex in 70% of individuals (30% were incorrect), while the chromosome-level Polar bear v1 HiC correctly determined the sex with 100% accuracy. When using the Dog assembly as RefGEN, we found 100% accuracy regardless of the number of mapped reads.

We also tested whether the two combinations of RefX and RefY used in each species data set (CowX/HumanY *vs* HumanX/HumanY for beluga; DogX/DogY *vs* HumanX/HumanY for polar bear) provided the same results. We observed a small fraction of contradictions in sex identification, where an individual was identified as a female when using one RefX/Y set, and as a male in the other RefX/Y set, despite the RefGEN and number of mapped reads being identical (Supplementary Table S5-S6). These contradictions only happen in bam files with < 5,000 reads, and they represent between 2.14% - 3.57% of all the sex identifications performed. When comparing sex identifications produced using identical RefGEN and number of mapped reads, but different RefX/Y combinations, results were identical in 94% of the pairwise comparisons (337 out of 360 comparisons, including both beluga and polar bear data sets). The inability to designate the sex of an individual with both combinations of RefX/Y and RefY was only observed in two comparison. In the remaining 6% of comparisons, 2% (eight comparisons) yielded contradicting sex identifications. In six of the comparisons the more distant HumanXY RefX/Y produced the correction results, in one comparison the DogXY gave the correct result (polar bear dataset) and, in the remaining comparison the CowXHumanY gave the correct result (beluga dataset) The last 4% (15 comparisons) comprised one determined sex (male or female) and one undetermined sex (X:A ratio of 0.7-0.8). We obtained contradicting sex determination only in comparisons using relatively few reads and with the low-quality Beluga v1 RefGEN (using 5,000 and 2,500 mapped reads), and with Beluga v3 and Polar bear v1 RefGEN (using 1,000 mapped reads).

## Discussion

Many biological specimens for which sex cannot be identified using morphology or other traditional approaches, such as faecal, environmental, and archaeological or palaeontological material, are also likely to contain highly contaminated and/or degraded DNA (Hrovatin & Kunej, 2018). SeXY was designed to utilise low-effort screening data for sex identification. Therefore, by assessing the reliability of SeXY to various levels of sequencing effort, we evaluate its applicability to such samples. Although our results differed between reference genomes, we show that less than 5,000 mapped reads can be used to accurately identify the biological sex of an individual, depending on the quality of the mapping reference. This finding opens a world of possibility for studies that employ low-effort shotgun sequencing approaches to identify specimens of sufficient preservation for deeper sequencing, but which discard any data/specimens not deemed of sufficient quality. By utilising our method, sequence information that would previously have been discarded can now be used to obtain sex-related evolutionary and biological insights. Although this has been done on several taxa (e.g., Gower et al., 2019; Pečnerová et al., 2017), our method, which does not require *a priori* sex-chromosome information from the target species or a reference panel of known females and males, will hopefully enable such analyses from a much wider range of species. Although only tested with up to 100,000 mapped reads, the increasing accuracy as the number of mapped reads increased means this method is also suitable for well-preserved specimens with more available sequencing data. In such cases, data could even be downsampled to increase computational speed.

### 1. Evaluation of synteny approach

SeXY identifies sex-linked scaffolds using a synteny approach (Grabherr et al., 2010), where the reference sex-chromosome assemblies (RefX and RefY) of a chromosome-level assembly from a closely related species is used to identify sequence similarities on the reference genome assembly (RefGEN). Although this method may have limitations due to computational time or the lacking identification of new (neo)-sex chromosomes (Marshall Graves, 2008; Nursyifa et al., 2021), our results show that SeXY could accurately determine the sex of the beluga and polar bear individuals, even with a relatively distant sex-chromosome assembly (in our case, human). In addition, the identification of sex-linked scaffolds is performed only once per reference genome assembly used, and hence computation time will not increase with the number of samples.

### 2. Number of mapped reads

Our finding of 100% accurate sex identification when mapping polar bear reads to the dog as RefGEN, even with only 1,000 mapped reads, was somewhat unexpected, as we anticipated a decline in mapping efficiency with increasing phylogenetic distance (Prasad et al., 2021). However, these results become less surprising when considering the mapping efficiency to each RefGEN. Although sex determination was 100% accurate down to 1,000 mapped reads when using these two species with ∼17% divergence, approximately four times as many raw reads are required to reach the target number of mapped reads, relative to when mapping to a conspecific RefGEN (Figure 2, Supplementary Table S4). Therefore, when < 5,000 endogenous reads are available, it is important to weigh the number of mapped reads *versus* the number of raw reads, to evaluate whether mapping to a conspecific reference genome or a phylogenetically distant reference genome is more beneficial. Although not tested here, alterations in mapping quality filters may facilitate the recovery of more mapped reads and thereby more accurate sex identification. However, decreased mapping quality may also result in misalignments, biasing results. Such low endogenous read counts are unlikely to arise when sequencing DNA from well-preserved samples, but it is much more common when considering highly degraded samples such as faecal, environmental, or subfossil material.

### 3. Quality and phylogenetic distance of the reference genome assembly

When comparing results produced by mapping beluga reads to the more fragmented Beluga v1 *versus* the more contiguous Beluga v3, we show the quality of the reference genome assembly can significantly impact the accuracy of sex determination. The two beluga assemblies are vastly different in quality, with scaffold N50s of 161 kb and 31 Mb, respectively. When considering < 50,000 mapped reads, the more fragmented Beluga v1 assembly could not be used to accurately determine sex. A fragmented reference genome assembly of lower quality, as with Beluga v1, may lead to difficulties in accurately identifying the sex-linked scaffolds, which our method is reliant on. Therefore, although not comprehensively investigated here, it is advisable to rather use a high-quality reference genome assembly from a phylogenetically more distant species, than a low-quality conspecific assembly. However, the accuracy of the X:A ratio using Beluga v1 as mapping reference provided 100% accuracy at 50,000 and above mapped reads. Therefore, we show that SeXY can still be used to accurately identify sex even if only a highly fragmented assembly is available, if the number of mapped reads is sufficiently high. This holds promise for the applicability of our method moving forward, as there are an increasing number of high-quality reference genomes available, and initiatives such as the Vertebrate Genome project aim to generate near error-free reference genome assemblies of many vertebrate species in the near future (Rhie et al., 2021).

Phylogenetic distance of the mapping reference genome assembly also appears to play a role. In the case of beluga mapped to the Orca RefGEN, comparisons using < 10,000 mapped reads were unable to accurately identify an individual’s sex. However, this finding may reflect the more fragmented assembly of the Orca (N50 = 13 Mb) relative to the other mapping references, as we were able to identify sex with 80% accuracy (89% excluding undetermined sex) using Cow as RefGEN down to 1,000 mapped reads. Furthermore, while Panda as RefGEN produced less consistent results for the polar bear than the two conspecific reference genome assemblies, the Panda results were far more consistent than when Orca was used as RefGEN for beluga, perhaps owing to the higher assembly quality of the Panda (N50 = 129 Mb). Thus, our results suggest that the quality of the reference genome assembly is far more important than phylogenetic distance between the species of interest and the mapping reference.

### 4. Recommendations and suggested guidelines

When relatively high numbers of reads are available (<50,000 mapped reads), our results show that both the fragmentation of the assembly and phylogenetic distance to the target species do not influence the accuracy of our method. Therefore, in cases such as these the choice of RefGen is at the discretion of the user.

However, for lower quality samples with fewer reads mapping, more discretion is required. Based on our results, the level of fragmentation of the RefGen is most important here. We found that a fragmented reference genome assembly, as with Beluga v1 (N50 161 kb), may lead to difficulties in accurately identifying the sex-linked scaffolds. Based on the clear impact of genome quality and a lack of clear impact of the phylogenetic distance between the target species and the mapping reference, we recommend using a more distant reference genome, if the quality of the closest reference genome is low and only relatively few mapped reads are available.

In conclusion, we demonstrate the method implemented in SeXY can accurately determine the sex of individuals based on very low sequencing effort, and when no conspecific chromosome-level assembly is available. The SeXY pipeline provides several advantages over previously implemented methods: SeXY (i) requires data from only a single individual (a mix of male and female individuals is not required), (ii) does not require assembled conspecific sex-chromosomes, or even a conspecific reference assembly, (ii) takes into account variation in coverage across the genome when calculating the X:A ratio, and (iv) can work on very low-coverage shotgun data, down to 1,000 mapped reads in many cases. Although we assessed the method based on XY sex chromosomes (as in mammals), the method can in theory be applied to any species with a heterogametic and a homogametic sex (e.g, birds, and some reptiles, fish, and insects), and in which the target and reference species share the same sex determination system.

## Acknowledgements

This work was supported by the Carlsberg Foundation Distinguished Associate Professor Fellowship (grant no CF16-0202) and the Villum Fonden Young Investigator program (grant no 13151) to E.D.L. A.A.C. was supported by a fellowship from Rubicon-NWO (project 019.183EN.005) and M.L. was supported by a fellowship from the Greenland Research Council.

## Data Availability Statement

All raw data used in this paper are publicly available in GenBank, NCBI and DNA Zoo. Accession codes can be found within the Materials and Methods and Supplementary Table S1-S3. SeXY pipeline is available on Github: https://github.com/andreidae/SeXY

## Author Contributions

A.A.C.: Conceptualization, Formal analysis, Writing -Original Draft, Visualisation; A.R-I: Conceptualization, Formal analysis, Writing - Review & Editing; M.L.: Conceptualization, Formal analysis, Writing - Review & Editing; M.S.: Conceptualization, Formal analysis, Writing - Review & Editing; M.V.W.: Conceptualization, Methodology, Writing - Review & Editing, Supervision; E.D.L.: Conceptualization, Resources, Writing - Review & Editing, Supervision, Funding acquisition.

## References

Aasen, E., & Medrano, J. F. (1990). Amplification of the ZFY and ZFX genes for sex identification in humans, cattle, sheep and goats. Bio/technology, 8(12), 1279–1281.

Ahlering, M. A., Hailer, F., Roberts, M. T., & Foley, C. (2011). A simple and accurate method to sex savannah, forest and Asian elephants using noninvasive sampling techniques. Molecular Ecology Resources, 11(5), 831–834.

Amos, B., Schlötterer, C., & Tautz, D. (1993). Social-structure of pilot whales revealed by analytical DNA profiling. Science, 260(5108), 670–672.

Bérubé, M., & Palsbøll, P. (1996). Identification of sex in cetaceans by multiplexing with three ZFX and ZFY specific primers. Molecular Ecology, 5(2), 283–287.

Bro-Jørgensen, M. H., Keighley, X., Ahlgren, H., Scharff-Olsen, C. H., Rosing-Asvid, A., Dietz, R., Ferguson, S. H., Gotfredsen, A. B., Jordan, P., Glykou, A., Lidén, K., & Olsen, M. T. (2021). Genomic sex identification of ancient pinnipeds using the dog genome. Journal of Archaeological Science, 127, 105321.

Bryja, J., & Konečný, A. (2003). Fast sex identification in wild mammals using PCR amplification of the Sry gene. Folia Zoo, 52(3), 269–274.

Buonasera, T., Eerkens, J., de Flamingh, A., Engbring, L., Yip, J., Li, H., Haas, R., DiGiuseppe, D., Grant, D., Salemi, M., Nijmeh, C., Arellano, M., Leventhal, A., Phinney, B., Byrd, B. F., Malhi, R. S., & Parker, G. (2020). A comparison of proteomic, genomic, and osteological methods of archaeological sex estimation. Scientific Reports, 10(1), 11897.

Bushnell, B. (2014). BBMap: A Fast, Accurate, Splice-Aware Aligner. No. LBNL-7065E. Conference: 9th Annual Genomics of Energy & Environment Meeting, Walnut Creek, CA, United States. https://www.semanticscholar.org/paper/f64dd54444a724574deb7710888091350eebb2b 9

Cabrera, A. A., Bérubé, M., Lopes, X. M., Louis, M., Oosting, T., Rey-Iglesia, A., Rivera-León, V. E., Székely, D., Lorenzen, E. D., & Palsbøll, P. J. (2021). A Genetic Perspective on Cetacean Evolution. Annual Review of Ecology, Evolution, and Systematics, 52, 131–151.

Curtis, C., Stewart, B. S., & Karl, S. A. (2007). Sexing pinnipeds with ZFX and ZFY loci. The Journal of Heredity, 98(3), 280–285.

de Flamingh, A., Coutu, A., Roca, A. L., & Malhi, R. S. (2020). Accurate Sex Identification of Ancient Elephant and Other Animal Remains Using Low-Coverage DNA Shotgun Sequencing Data. G3, 10(4), 1427–1432.

Dobson, F. S., & Stephen Dobson, F. (1982). Competition for mates and predominant juvenile male dispersal in mammals. In Animal Behaviour (Vol. 30, Issue 4, pp. 1183– 1192). https://doi.org/10.1016/s0003-3472(82)80209-1

Dudchenko, O., Batra, S. S., Omer, A. D., Nyquist, S. K., Hoeger, M., Durand, N. C., Shamim, M. S., Machol, I., Lander, E. S., Aiden, A. P., & Aiden, E. L. (2017). De novo assembly of the Aedes aegypti genome using Hi-C yields chromosome-length scaffolds. Science, 356(6333), 92–95.

Dudchenko, O., Shamim, M. S., Batra, S. S., Durand, N. C., Musial, N. T., Mostofa, R., Pham, M., St Hilaire, B. G., Yao, W., Stamenova, E., Hoeger, M., Nyquist, S. K., Korchina, V., Pletch, K., Flanagan, J. P., Tomaszewicz, A., McAloose, D., Estrada, C. P., Novak, B. J., … Aiden, E. L. (2018). The Juicebox Assembly Tools module facilitates de novo assembly of mammalian genomes with chromosome-length scaffolds for under $1000. In bioRxiv (p. 254797). https://doi.org/10.1101/254797

Durnin, M. E., Palsbøll, P. J., Ryder, O. A., & McCullough, D. R. (2007). A reliable genetic technique for sex determination of giant panda (Ailuropoda melanoleuca) from non-invasively collected hair samples. Conservation Genetics, 8(3), 715–720.

Ellegren, H. (2000). Evolution of the avian sex chromosomes and their role in sex determination. In Trends in Ecology & Evolution (Vol. 15, Issue 5, pp. 188–192). https://doi.org/10.1016/s0169-5347(00)01821-8

Fairbairn, D. J., Blanckenhorn, W. U., & Székely, T. (2008). Sex, Size and Gender Roles: Evolutionary Studies of Sexual Size Dimorphism. OUP Oxford.

Fan, H., Wu, Q., Wei, F., Yang, F., Ng, B. L., & Hu, Y. (2019). Chromosome-level genome assembly for giant panda provides novel insights into Carnivora chromosome evolution. Genome Biology, 20(1), 267.

Foote, A. D., Liu, Y., Thomas, G. W. C., Vinař, T., Alföldi, J., Deng, J., Dugan, S., van Elk, C. E., Hunter, M. E., Joshi, V., Khan, Z., Kovar, C., Lee, S. L., Lindblad-Toh, K., Mancia, A., Nielsen, R., Qin, X., Qu, J., Raney, B. J., … Gibbs, R. A. (2015). Convergent evolution of the genomes of marine mammals. Nature Genetics, 47(3), 272–275.

Gower, G., Fenderson, L. E., Salis, A. T., Helgen, K. M., van Loenen, A. L., Heiniger, H., Hofman-Kamińska, E., Kowalczyk, R., Mitchell, K. J., Llamas, B., & Cooper, A. (2019). Widespread male sex bias in mammal fossil and museum collections. Proceedings of the National Academy of Sciences of the United States of America, 116(38), 19019– 19024.

Grabherr, M. G., Russell, P., Meyer, M., Mauceli, E., Alföldi, J., Di Palma, F., & Lindblad-Toh, K. (2010). Genome-wide synteny through highly sensitive sequence alignment: Satsuma. Bioinformatics, 26(9), 1145–1151.

Greenwood, P. J. (1980). Mating systems, philopatry and dispersal in birds and mammals. In Animal Behaviour (Vol. 28, Issue 4, pp. 1140–1162). https://doi.org/10.1016/s0003-3472(80)80103-5

Hrovatin, K., & Kunej, T. (2018). Genetic sex determination assays in 53 mammalian species: Literature analysis and guidelines for reporting standardization. Ecology and Evolution, 8(2), 1009–1018.

Hu, Y., Wu, Q., Ma, S., Ma, T., Shan, L., Wang, X., Nie, Y., Ning, Z., Yan, L., Xiu, Y., & Wei, F. (2017). Comparative genomics reveals convergent evolution between the bamboo-eating giant and red pandas. Proceedings of the National Academy of Sciences of the United States of America, 114(5), 1081–1086.

íKongsstovu, S., Dahl, H. A., Gislason, H., Homrum, E., Jacobsen, J. A., Flicek, P., & Mikalsen, S.-O. (2020). Identification of male heterogametic sex-determining regions on the Atlantic herring Clupea harengus genome. Journal of Fish Biology, 97(1), 190–201.

Jones, S. J. M., Taylor, G. A., Chan, S., Warren, R. L., Hammond, S. A., Bilobram, S., Mordecai, G., Suttle, C. A., Miller, K. M., Schulze, A., Chan, A. M., Jones, S. J., Tse, K., Li, I., Cheung, D., Mungall, K. L., Choo, C., Ally, A., Dhalla, N., … Haulena, M. (2017). The Genome of the Beluga Whale (Delphinapterus leucas). Genes, 8(12). https://doi.org/10.3390/genes8120378

Kirch, M., Romundset, A., Gilbert, M. T. P., Jones, F. C., & Foote, A. D. (2021). Ancient and modern stickleback genomes reveal the demographic constraints on adaptation. Current Biology: CB, 31(9), 2027–2036.e8.

Kumar, S., Stecher, G., Suleski, M., & Hedges, S. B. (2017). TimeTree: A Resource for Timelines, Timetrees, and Divergence Times. Molecular Biology and Evolution, 34(7), 1812–1819.

Li, H. (2013). Aligning sequence reads, clone sequences and assembly contigs with BWA-MEM. arXiv E-Prints, arXiv:1303.3997.

Li, H., Handsaker, B., Wysoker, A., Fennell, T., Ruan, J., Homer, N., Marth, G., Abecasis, G., Durbin, R., & 1000 Genome Project Data Processing Subgroup. (2009). The Sequence Alignment/Map format and SAMtools. In Bioinformatics (Vol. 25, Issue 16, pp. 2078–2079). https://doi.org/10.1093/bioinformatics/btp352

Lindblad-Toh, K., Wade, C. M., Mikkelsen, T. S., Karlsson, E. K., Jaffe, D. B., Kamal, M., Clamp, M., Chang, J. L., Kulbokas, E. J., 3rd, Zody, M. C., Mauceli, E., Xie, X., Breen, M., Wayne, R. K., Ostrander, E. A., Ponting, C. P., Galibert, F., Smith, D. R., DeJong, P. J., … Lander, E. S. (2005). Genome sequence, comparative analysis and haplotype structure of the domestic dog. Nature, 438(7069), 803–819.

Liu, S., Lorenzen, E. D., Fumagalli, M., Li, B., Harris, K., Xiong, Z., Zhou, L., Korneliussen, T. S., Somel, M., Babbitt, C., Wray, G., Li, J., He, W., Wang, Z., Fu, W., Xiang, X., Morgan, C. C., Doherty, A., O’Connell, M. J., … Wang, J. (2014). Population genomics reveal recent speciation and rapid evolutionary adaptation in polar bears. Cell, 157(4), 785– 794.

Louis, M., Skovrind, M., Garde, E., Heide-Jørgensen, M. P., Szpak, P., & Lorenzen, E. D. (2021). Population-specific sex and size variation in long-term foraging ecology of belugas and narwhals. Royal Society Open Science, 8(2), 202226.

Malisa, A., Gwakisa, P., Balthazary, S., Wasser, S., & Mutayoba, B. (2005). Species and gender differentiation between and among domestic and wild animals using mitochondrial and sex-linked DNA markers. African Journal of Biotechnology, 4(11). https://doi.org/10.4314/ajb.v4i11.71391

Marshall Graves, J. A. (2008). Weird animal genomes and the evolution of vertebrate sex and sex chromosomes. Annual Review of Genetics, 42, 565–586.

McGowen, M. R., Tsagkogeorga, G., Álvarez-Carretero, S., Dos Reis, M., Struebig, M., Deaville, R., Jepson, P. D., Jarman, S., Polanowski, A., Morin, P. A., & Rossiter, S. J. (2020). Phylogenomic Resolution of the Cetacean Tree of Life Using Target Sequence Capture. Systematic Biology, 69(3), 479–501.

Mittnik, A., Wang, C.-C., Svoboda, J., & Krause, J. (2016). A Molecular Approach to the Sexing of the Triple Burial at the Upper Paleolithic Site of Dolní Věstonice. PloS One, 11(10), e0163019.

Moore, C. R. (1925). Sex determination and sex differentiation in birds and mammals. In The American Naturalist (Vol. 59, Issue 661, pp. 177–189). https://doi.org/10.1086/280026

Nistelberger, H. M., Pálsdóttir, A. H., Star, B., Leifsson, R., Gondek, A. T., Orlando, L., Barrett, J. H., Hallsson, J. H., & Boessenkool, S. (2019). Sexing Viking Age horses from burial and non-burial sites in Iceland using ancient DNA. In Journal of Archaeological Science (Vol. 101, pp. 115–122). https://doi.org/10.1016/j.jas.2018.11.007

Nursyifa, C., Brüniche-Olsen, A., Erill, G. G., Heller, R., & Albrechtsen, A. (2021). Joint identification of sex and sex-linked scaffolds in non-model organisms using low depth sequencing data. Molecular Ecology Resources, 00, 1–10.

Palmer, D. H., Rogers, T. F., Dean, R., & Wright, A. E. (2019). How to identify sex chromosomes and their turnover. Molecular Ecology, 28(21), 4709–4724.

Pečnerová, P., Díez-Del-Molino, D., Dussex, N., Feuerborn, T., von Seth, J., van der Plicht, J., Nikolskiy, P., Tikhonov, A., Vartanyan, S., & Dalén, L. (2017). Genome-based sexing provides clues about behavior and social structure in the woolly mammoth. Current Biology: CB, 27(22), 3505–3510.e3.

Pomp, D., Good, B. A., Geisert, R. D., Corbin, C. J., & Conley, A. J. (1995). Sex identification in mammals with polymerase chain reaction and its use to examine sex effects on diameter of day-10 or -11 pig embryos. Journal of Animal Science, 73(5), 1408–1415.

Prasad, A., Lorenzen, E. D., & Westbury, M. V. (2021). Evaluating the role of reference-genome phylogenetic distance on evolutionary inference. Molecular Ecology Resources. https://doi.org/10.1111/1755-0998.13457

Quinlan, A. R., & Hall, I. M. (2010). BEDTools: a flexible suite of utilities for comparing genomic features. In Bioinformatics (Vol. 26, Issue 6, pp. 841–842). https://doi.org/10.1093/bioinformatics/btq033

Rhie, A., McCarthy, S. A., Fedrigo, O., Damas, J., Formenti, G., Koren, S., Uliano-Silva, M., Chow, W., Fungtammasan, A., Kim, J., Lee, C., Ko, B. J., Chaisson, M., Gedman, G. L., Cantin, L. J., Thibaud-Nissen, F., Haggerty, L., Bista, I., Smith, M., … Jarvis, E. D. (2021). Towards complete and error-free genome assemblies of all vertebrate species. Nature, 592(7856), 737–746.

Schubert, M., Ermini, L., Der Sarkissian, C., Jónsson, H., Ginolhac, A., Schaefer, R., Martin, M. D., Fernández, R., Kircher, M., McCue, M., Willerslev, E., & Orlando, L. (2014). Characterization of ancient and modern genomes by SNP detection and phylogenomic and metagenomic analysis using PALEOMIX. Nature Protocols, 9(5), 1056–1082.

Schubert, M., Lindgreen, S., & Orlando, L. (2016). AdapterRemoval v2: rapid adapter trimming, identification, and read merging. BMC Research Notes, 9, 88.

Sinding, M.-H. S., Tervo, O. M., Grønnow, B., Gulløv, H. C., Toft, P. A., Bachmann, L., Fietz, K., Rekdal, S. L., Christoffersen, M. F., Heide-Jørgensen, M. P., Olsen, M. T., & Foote, A. D. (2016). Sex determination of baleen whale artefacts: Implications for ancient DNA use in zooarchaeology. Journal of Archaeological Science: Reports, 10, 345–349.

Skoglund, P., Storå, J., Götherström, A., & Jakobsson, M. (2013). Accurate sex identification of ancient human remains using DNA shotgun sequencing. Journal of Archaeological Science, 40(12), 4477–4482.

Skovrind, M., Castruita, J. A. S., Haile, J., Treadaway, E. C., Gopalakrishnan, S., Westbury, M. V., Heide-Jørgensen, M. P., Szpak, P., & Lorenzen, E. D. (2019). Hybridization between two high Arctic cetaceans confirmed by genomic analysis. In Scientific Reports (Vol. 9, Issue 1). https://doi.org/10.1038/s41598-019-44038-0

Westbury, M. V., Petersen, B., Garde, E., Heide-Jørgensen, M. P., & Lorenzen, E. D. (2019). Narwhal Genome Reveals Long-Term Low Genetic Diversity despite Current Large Abundance Size. iScience. https://doi.org/10.1016/j.isci.2019.03.023

Zimin, A. V., Delcher, A. L., Florea, L., Kelley, D. R., Schatz, M. C., Puiu, D., Hanrahan, F., Pertea, G., Van Tassell, C. P., Sonstegard, T. S., Marçais, G., Roberts, M., Subramanian, P., Yorke, J. A., & Salzberg, S. L. (2009). A whole-genome assembly of the domestic cow, Bos taurus. Genome Biology, 10(4), R42.

