## Supplementary for "How low can you go: sex identification from low-quantity sequencing data despite lacking assembled sex chromosomes"

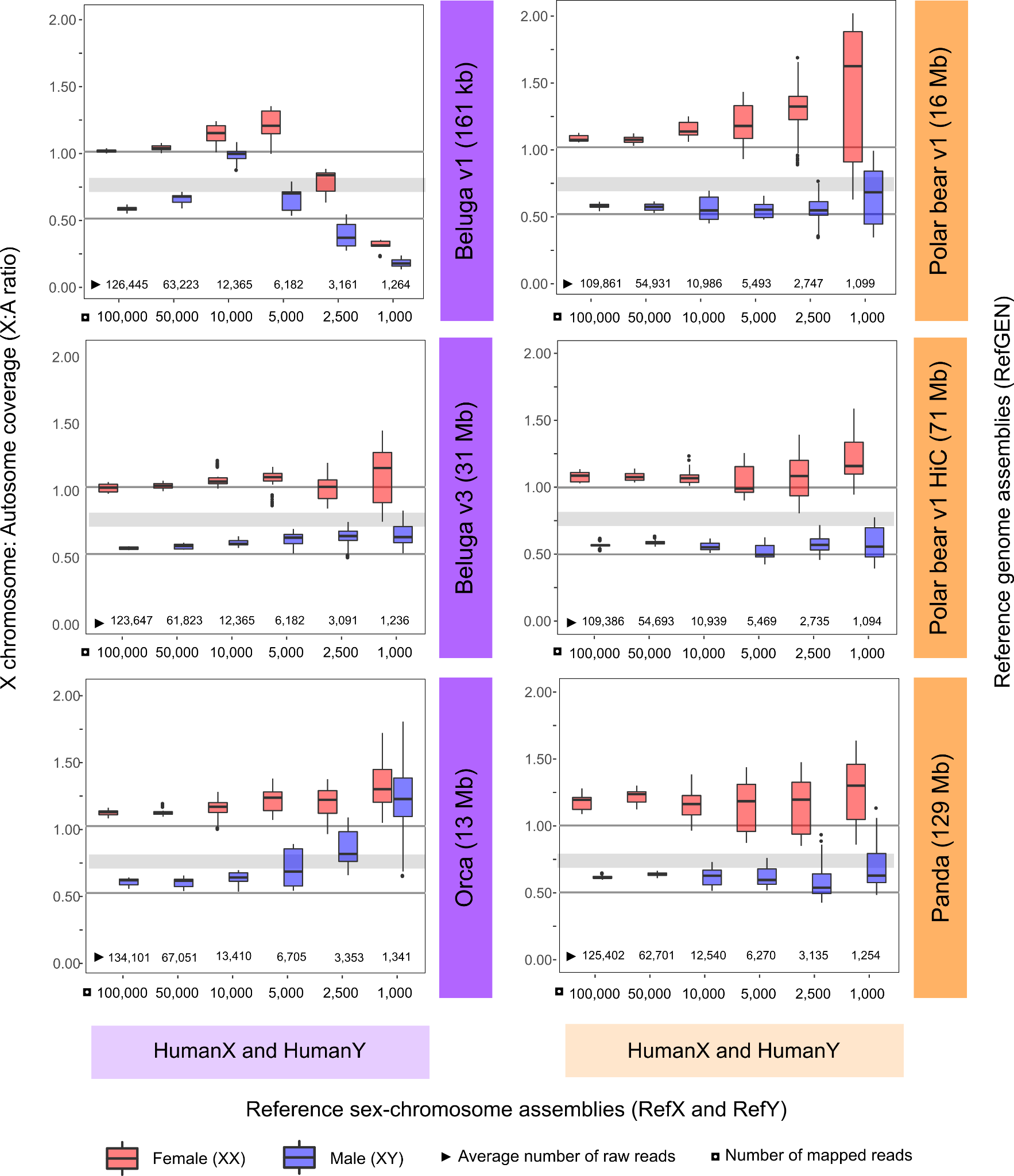

**Supplementary Figure S1. Sex determination of beluga and polar bear individuals using four reference genome assemblies (RefGEN), HumanX and HumanY as reference sex-chromosome assembly (RefX and RefY), and various numbers of mapped reads.** The ten beluga and ten polar bear individuals tested both comprised five females (red) and five males (blue). X axis shows number of mapped reads (square) and average number of raw reads necessary to obtain the required number of mapped reads (triangle). Y axis shows comparison of X chromosome and autosome coverage (X:A ratio) for each combination of RefGEN, RefX and RefY, and number of mapped reads. Individuals were determined as females if their X:A ratio was >=0.8, and as males if their X:A ratio was <=0.7. Grey shaded horizontal bars indicate an X:A ratio of 0.7-0.8, which we interpreted as undetermined sex.

**Supplementary Table S1. List, accession numbers, and sex of the ten beluga and ten polar bear samples used in this study to assess the SeXY pipeline.**

| **Species** | **Sample ID** | **Biosample** | **SRA** | **Sex** |
| --- | --- | --- | --- | --- |
| Beluga | DL_CIB01 | SAMN12723975 | SRS5486914 | Female |
| Beluga | DL_CIB03 | SAMN12724024 | SRS5486912 | Female |
| Beluga | DL_CIB04 | SAMN12724025 | SRS5486911 | Female |
| Beluga | DL_CIB06 | SAMN12724052 | SRS5486910 | Female |
| Beluga | DL_CIB11 | SAMN12724179 | SRS5486906 | Female |
| Beluga | BB_1402 | SAMN12724224 | SRS5486901 | Male |
| Beluga | DL_CIB02 | SAMN12724023 | SRS5486913 | Male |
| Beluga | DL_CIB07 | SAMN12724098 | SRS5486909 | Male |
| Beluga | DL_CIB08 | SAMN12724105 | SRS5486908 | Male |
| Beluga | DL_CIB10 | SAMN12724146 | SRS5486907 | Male |
| Polar bear | PB_2 | SAMN02261850 | SRR942245 | Female |
| Polar bear | PB_49 | SAMN02261863 | SRR942258 | Female |
| Polar bear | PB_54 | SAMN02261844 | SRR942239 | Female |
| Polar bear | PB_69 | SAMN02261862 | SRR942257 | Female |
| Polar bear | PB_8 | SAMN02261849 | SRR942244 | Female |
| Polar bear | PB_37 | SAMN02261806 | SRR942198 | Male |
| Polar bear | PB_38 | SAMN02261807 | SRR942199 | Male |
| Polar bear | PB_50 | SAMN02261802 | SRR942194 | Male |
| Polar bear | PB_52 | SAMN02261803 | SRR942195 | Male |
| Polar bear | PB_53 | SAMN02261804 | SRR942196 | Male |

**Supplementary Table S2. Overview of reference genome assemblies (RefGEN) used for mapping of target species reads.**

| **Species** | **MappingID** | **Biosample** | **Genome assembly accession** | **Assembly level** | **Sex** | **N50 (bp)** |
| --- | --- | --- | --- | --- | --- | --- |
| Beluga | Beluga v1 | SAMN06216270 | GCA_002288925.1 | Contig | Female | 161,467 |
| Beluga | Beluga v3 | SAMN06216270 | GCA_002288925.3 | Scaffold | Female | 31,183,418 |
| Orca | Orca | SAMN01180276 | GCA_000331955.2 | Scaffold | Female | 12,735,091 |
| Cow | Cow | SAMN03145444 | GCA_002263795.2 | Chromosome | Female | 103,308,737 |
| Polar bear | Polar bear v1 | SAMN02729226 | GCA_000687225.1 | Scaffold | Male | 15,940,661 |
| Polar bear | Polar bear v1 HiC | SAMN02729226 | UrsMar_1.0_HiC | Chromosome draft | Male | 71,276,975 |
| Dog | Dog | SAMN14478636 | GCA_014441545.1 | Chromosome | Male | 64,037,277 |
| Panda | Panda | SAMN04193337 | GCA_002007445.2 | Chromosome | Female | 129,245,720 |

**Supplementary Table S3. List of reference sex-chromosome assemblies (RefX and RefY) used by satsuma synteny v2.1 to identify sex-linked scaffolds in each reference genome assembly (RefGEN).**

| **Species** | **NCBI reference** | **Sex-chromosome**  **assembly accession** | **Sex chromosome** |
| --- | --- | --- | --- |
| Cow | NC_037357.1 | GCA_002263795.2 | X |
| Dog | NC_051843.1 | GCA_014441545.1 | X |
| Dog | NC_051844.1 | GCA_014441545.1 | Y |
| Human | NC_000023.11 | GCA_000001405.28 | X |
| Human | NC_000024.10 | GCA_000001405.28 | Y |

**Supplementary Table S4. Mapping statistics for each beluga and polar bear individual mapped to the various reference genome assemblies (RefGEN) from the Paleomix summary files.**  To reduce computational time during the mapping step, all read files were randomly downsampled to one million reads prior to mapping**.**

See table as spreadsheet

**Supplementary Table S5. X:A ratio estimates across samples for the different reference genome (RefGEN) and sex-chromosome (RefX, RefY) assemblies**

See table as spreadsheet

**Supplementary Table S6. Number of correctly, undetermined, and incorrectly determined sex for ten beluga and ten polar bear individuals of known sex across tested combinations of reference genome assembly (RefGEN), reference sex-chromosome assembly (RefX and RefY), and numbers of mapped reads.** Sex determination of each indvidual is calculated using the average value of ten replicates, assuming a threshold of <= 0.7: male, 0.7-0.8: undetermined sex; >= 0.8: female.

|  | **Species** | **Beluga** | | | | | | | **Polar bear** | | | | | | |
| --- | --- | --- | --- | --- | --- | --- | --- | --- | --- | --- | --- | --- | --- | --- | --- |
| Number of mapped reads | **RefGen** | **Beluga v1** | | **Beluga v3** | | **Orca** | | **Cow** | **Polar bear v1** | | **Polar bear v1 HiC** | | **Panda** | | **Dog** |
|  | **RefX and RefY** | **CowX and HumanY** | **HumanX and HumanY** | **CowX and HumanY** | **HumanX and HumanY** | **CowX and HumanY** | **HumanX and HumanY** | **CowX and HumanY** | **DogX and DogY** | **HumanX and HumanY** | **DogX and DogY** | **HumanX and HumanY** | **DogX and DogY** | **HumanX and HumanY** | **DogX and DogY** |
| 100,000 | Correct | 10 | 10 | 10 | 10 | 10 | 10 | 10 | 10 | 10 | 10 | 10 | 10 | 10 | 10 |
|  | Undetermined | 0 | 0 | 0 | 0 | 0 | 0 | 0 | 0 | 0 | 0 | 0 | 0 | 0 | 0 |
|  | Incorrect | 0 | 0 | 0 | 0 | 0 | 0 | 0 | 0 | 0 | 0 | 0 | 0 | 0 | 0 |
| 50,000 | Correct | 10 | 10 | 10 | 10 | 10 | 10 | 10 | 10 | 10 | 10 | 10 | 10 | 10 | 10 |
|  | Undetermined | 0 | 0 | 0 | 0 | 0 | 0 | 0 | 0 | 0 | 0 | 0 | 0 | 0 | 0 |
|  | Incorrect | 0 | 0 | 0 | 0 | 0 | 0 | 0 | 0 | 0 | 0 | 0 | 0 | 0 | 0 |
| 10,000 | Correct | 5 | 5 | 10 | 10 | 9 | 10 | 10 | 10 | 10 | 10 | 10 | 10 | 10 | 10 |
|  | Undetermined | 0 | 0 | 0 | 0 | 1 | 0 | 0 | 0 | 0 | 0 | 0 | 0 | 0 | 0 |
|  | Incorrect | 5 | 5 | 0 | 0 | 0 | 0 | 0 | 0 | 0 | 0 | 0 | 0 | 0 | 0 |
| 5,000 | Correct | 5 | 9 | 9 | 10 | 8 | 8 | 10 | 10 | 10 | 10 | 10 | 10 | 10 | 10 |
|  | Undetermined | 1 | 1 | 1 | 0 | 1 | 0 | 0 | 0 | 0 | 0 | 0 | 0 | 0 | 0 |
|  | Incorrect | 4 | 0 | 0 | 0 | 1 | 2 | 0 | 0 | 0 | 0 | 0 | 0 | 0 | 0 |
| 2,500 | Correct | 10 | 8 | 10 | 9 | 5 | 6 | 8 | 10 | 10 | 10 | 10 | 9 | 9 | 10 |
|  | Undetermined | 0 | 1 | 0 | 1 | 2 | 2 | 2 | 0 | 0 | 0 | 0 | 1 | 0 | 0 |
|  | Incorrect | 0 | 1 | 0 | 0 | 3 | 2 | 0 | 0 | 0 | 0 | 0 | 0 | 1 | 0 |
| 1,000 | Correct | 5 | 5 | 8 | 9 | 6 | 6 | 8 | 7 | 7 | 10 | 8 | 8 | 8 | 10 |
|  | Undetermined | 0 | 0 | 2 | 1 | 0 | 0 | 1 | 0 | 1 | 0 | 2 | 1 | 1 | 0 |
|  | Incorrect | 5 | 5 | 0 | 0 | 4 | 4 | 1 | 3 | 2 | 0 | 0 | 1 | 1 | 0 |
| Total |  | 60 | 60 | 60 | 60 | 60 | 60 | 60 | 60 | 60 | 60 | 60 | 60 | 60 | 60 |

**Supplementary Table S7: Summary table showing percentage of correct sex determination across tested combinations of reference genome assembly (RefGEN), reference sex-chromosome assembly (RefX and RefY), and number of mapped reads.** Results are shown for the beluga data and the cetacean/cow RefGEN assemblies tested (left columns), and for the polar bear data and the bear/dog RefGEN assemblies tested (right columns). The value below each RefGEN indicates the assembly N50. For cells with two estimates, the left value indicates estimates including undetermined sex, and the right value indicates estimates excluding undetermined sex. Only one value is included if both estimates were the same. Percentages in each cell are based on ten sampled individuals; five females and five males. Sex determination for each individual was calculated using the average value of ten replicates, assuming a threshold of <= 0.7: male; 0.7-0.8: undetermined sex; >= 0.8: female.

|  | **Beluga** | | | | **Polar bear** | | | |
| --- | --- | --- | --- | --- | --- | --- | --- | --- |
| **Number of mapped reads** | **Beluga v1** | **Beluga v3** | **Orca** | **Cow** | **Polar bear v1** | **Polar bear v1 HiC** | **Panda** | **Dog** |
|  | **161 kb** | **31 Mb** | **13 Mb** | **103 Mb** | **16 Mb** | **71 Mb** | **129 Mb** | **64 Mb** |
|  | **HumanX and HumanY** | | | | | | | |
| 100,000 | 100 | 100 | 100 | - | 100 | 100 | 100 | - |
| 50,000 | 100 | 100 | 100 | - | 100 | 100 | 100 | - |
| 10,000 | 50 | 100 | 100 | - | 100 | 100 | 100 | - |
| 5,000 | 90/100 | 100 | 80 | - | 100 | 100 | 100 | - |
| 2,500 | 80/89 | 90/100 | 60/75 | - | 100 | 100 | 90 | - |
| 1,000 | 50 | 90/100 | 60 | - | 70/78 | 80/100 | 80/89 | - |
